# Apoptosis is not conserved in plants as revealed by critical examination of a model for plant apoptosis-like cell death

**DOI:** 10.1101/2020.09.26.314583

**Authors:** Elena A. Minina, Adrian N. Dauphinee, Florentine Ballhaus, Vladimir Gogvadze, Andrei P. Smertenko, Peter V. Bozhkov

**Author notes:** Correspondence: EAM, PVB. equal contribution (Elena A. Minina, Adrian N. Dauphinee).

## Abstract

**Background:** Animals and plants diverged over one billion years ago and evolved unique mechanisms for many cellular processes, including cell death. One of the most well-studied cell-death programmes in animals, apoptosis, involves gradual cell dismantling and engulfment of cellular fragments, apoptotic bodies, through phagocytosis. However, rigid cell walls prevent plant cell fragmentation and thus apoptosis is not applicable for executing cell death in plants. Furthermore, plants are devoid of the key components of apoptotic machinery, including phagocytosis as well as caspases and Bcl-2 family proteins. Nevertheless, the concept of plant “apoptosis-like programmed cell death” (AL-PCD) is widespread. This is largely due to superficial morphological resemblances between plant cell death and apoptosis; in particular between protoplast shrinkage in plant cells killed by various stimuli and animal cell volume decrease preceding fragmentation into apoptotic bodies.

**Results:** Here, we provide a comprehensive spatio-temporal analysis of cytological and biochemical events occurring in plant cells subjected to heat shock at 40-55°C and 85°C, the experimental conditions typically used to trigger AL-PCD and necrotic cell death, respectively. We show that cell death under both conditions was not accompanied by membrane blebbing or formation of apoptotic bodies, as would be expected during apoptosis. Instead, we observed instant and irreversible permeabilization of the plasma membrane and ATP depletion. These processes did not depend on mitochondrial functionality or the presence of Ca^2+^ and could not be prevented by an inhibitor of ferroptosis. We further reveal that the lack of protoplast shrinkage at 85°C, the only striking morphological difference between cell deaths induced by 40-55°C or 85°C heat shock, is a consequence of the fixative effect of the high temperature on intracellular contents.

**Conclusions:** We conclude that heat shock-induced cell death is an energy-independent process best matching definition of necrosis. Although the initial steps of this necrotic cell death could be genetically regulated, classifying it as apoptosis or AL-PCD is a terminological misnomer. Our work supports the viewpoint that apoptosis is not conserved across animal and plant kingdoms and demonstrates the importance of focusing on plant-specific aspects of cell death pathways.

## Background

Upon acute stress, cells might succumb to an accidental necrosis, which is essentially an unregulated collapse of cellular functions. In contrast, regulated cell death (RCD) is a genetically encoded process of cellular suicide, which is built into physiology of a multicellular organism (1). For example, programmed cell death (PCD), a subcategory of RCD intimately associated with normal development, counteracts cell proliferation by removing aged cells and plays an essential role in morphogenesis by eliminating surplus cells and shaping new structures (2–5). Other examples of RCD are implicated in stress responses, restricting the spread of pathogens through tissues and confining damage caused by abiotic factors (5–7). For historical reasons, in plant research field RCD is typically referred to as PCD. However, to enable accurate comparison of cell death in animal and plant kingdoms, we apply here the more inclusive term RCD to plants as well.

The term apoptosis (ancient Greek άπόπτωσις, apóptōsis, “falling off”) was coined in the pivotal work of John Kerr and colleagues (8) to name the newly described type of animal RCD after the death of leaves programmed to occur in many plant species at the end of each growth season. Since then, the apoptotic cell death became one of the most studied pathways in animal model systems (for review see (9)). Understanding the plant RCD machinery advanced much slower, and progress in this direction unfortunately suffers from attempts to extrapolate the knowledge obtained on animal models.

The utmost importance of RCD for all multicellular organisms strongly suggests that this process should be evolutionary conserved. Indeed, RCD in animals and plants share quite a number of conceptual similarities that can be attributed to the common features of all eukaryotic cells, e.g. dependency on ATP and protein synthesis, regulated proteolysis, and often implication of a careful dismantling of a dying cell so as to protect neighbouring cells from its toxic content (10,11). However, RCD is an integral part of organismal developmental program and a key player in adaptation to environmental conditions, response to stress stimuli and pathogens, all of which would greatly vary depending on the life strategy of an organism. Thus, differences in cell organization, motility, genome maintenance, nutrition and body plan plasticity between animals and plants (12,13) could not have been shaped without tailoring their corresponding RCD machineries.

One of the most apparent differences between animals and plants is cell motility. Unlike animal cells, rigid carbohydrate walls of plant cells prevent guided migration within a multicellular organism (14). This fact alone illustrates the profound differences that can be expected between plant and animal RCD. Namely, due to the cellular immobility, plants lack mechanisms for preventing spread of malignant cells and do not have metastatic cancer. Thus, somatic mutations in plant and animal cells pose fundamentally different level of danger to their organisms, which shaped distinct strategies of the RCD regulation by the DNA-damage controlling machinery (15,16). Furthermore, execution of the RCD should also be different, as the cell walls make animal strategies of cellular disassembly not applicable for plant cells. For instance, apoptotic cells undergo fragmentation into smaller apoptotic bodies that can be taken up by phagocytes in a highly regulated fashion involving decoration of the apoptotic bodies with phosphatidylserine (17). The lack of phagocytes in plants means that apoptotic bodies would remain trapped within the cell wall cage, eventually undergoing decay and spilling toxic contents on the neighbouring cells. Thus, plant RCD should rely on specific regulation and execution mechanisms (11).

Although apoptosis was initially considered a synonym for RCD or PCD, numerous non-apoptotic types of RCD have been discovered in animals in the past two decades (1,18,19), such as paraptosis (20), several forms of regulated necrosis (necroptosis (21), ferroptosis (22), pyroptosis (23)), and others. Since animals possess multiple RCD pathways shaped to best fit the physiology of the organism under varying conditions, it is safe to assume that plant evolution shaped plant-specific RCD mechanisms.

The continuous search for biochemical manifestations of apoptosis in plants thus far has yielded unconvincing results. For instance, plants lack the core apoptotic regulators caspases and B-cell lymphoma 2 (Bcl-2) family proteins (24,25). Furthermore, although increase in caspase-like proteolytic activity has been detected during plant cell death (26), the analysis of the plant genome sequence data demonstrated lack of caspases (27). It remains to be further determined what subset of plant proteases is responsible for cell death-associated caspase-like activity, what are their substrates and what is the mechanistic role of limited proteolysis in the activation and execution of RCD (for review see (28)). The closest plant structural homologues of caspases, metacaspases, are indeed implicated in plant RCD (29,30), however, their substrate specificity and regulation are profoundly different from caspases (31,32). These facts point out the lack of instruments for apoptotic cell death in plants and support the existence of plant-specific RCD mechanisms.

Another example is Bcl-2 family proteins that play either pro- or anti-apoptotic functions in animals by regulating permeabilization of the outer mitochondrial membrane followed by the release of cytochrome *c* (33). It has been shown that ectopic expression of animal genes encoding pro- or anti-apoptotic Bcl-2 family members in plants can enhance or suppress cell death, respectively (34,35). However, it is not surprising that animal proteins inducing permeabilization of the outer mitochondrial membrane would cause mitochondrial dysfunction eventually leading to cell death in heterologous systems. Furthermore, although interaction of animal anti-apoptotic Bcl-2 protein with as yet to be identified plant protein might indeed decrease plant cell susceptibility to death stimuli, the lack of Bcl-2 genes in plant genomes points out that plant RCD does not rely on activity of these proteins and is thus regulated differently from the apoptotic pathway (36,37). Nevertheless, original research and reviews on apoptotic hallmarks in plant cell death are still being published (for recent reviews, see (38–41)), propagating controversial conclusions that require careful consideration.

Apoptotic cell death has typical hallmarks: (i) it is an active, ATP- and caspase-dependent process (42,43); (ii) the plasma membrane (PM) integrity is retained throughout the cell death process and phosphatidylserine is exposed on the outer membrane surface as an “eat-me” signal for phagocytes (44,45); (iii) cell shrinkage (or apoptotic volume decrease, AVD) is followed by chromatin condensation and nuclear segmentation (1,18,46); (iv) PM blebbing results in cell fragmentation into apoptotic bodies (10,47) and their subsequent phagocytosis (17).

The term “apoptosis-like PCD” (AL-PCD) was coined due to the reported morphological and biochemical similarities between animal apoptosis and stress-associated plant cell death, primarily AVD-like protoplast shrinkage, but also Ca^2+^- and ATP-dependency, maintenance of PM integrity and involvement of caspase-like activity (48–51). However, studies reporting on such similarities contain a number of controversies that need to be addressed before drawing conclusions (**Table 1**).

**Table 1.**
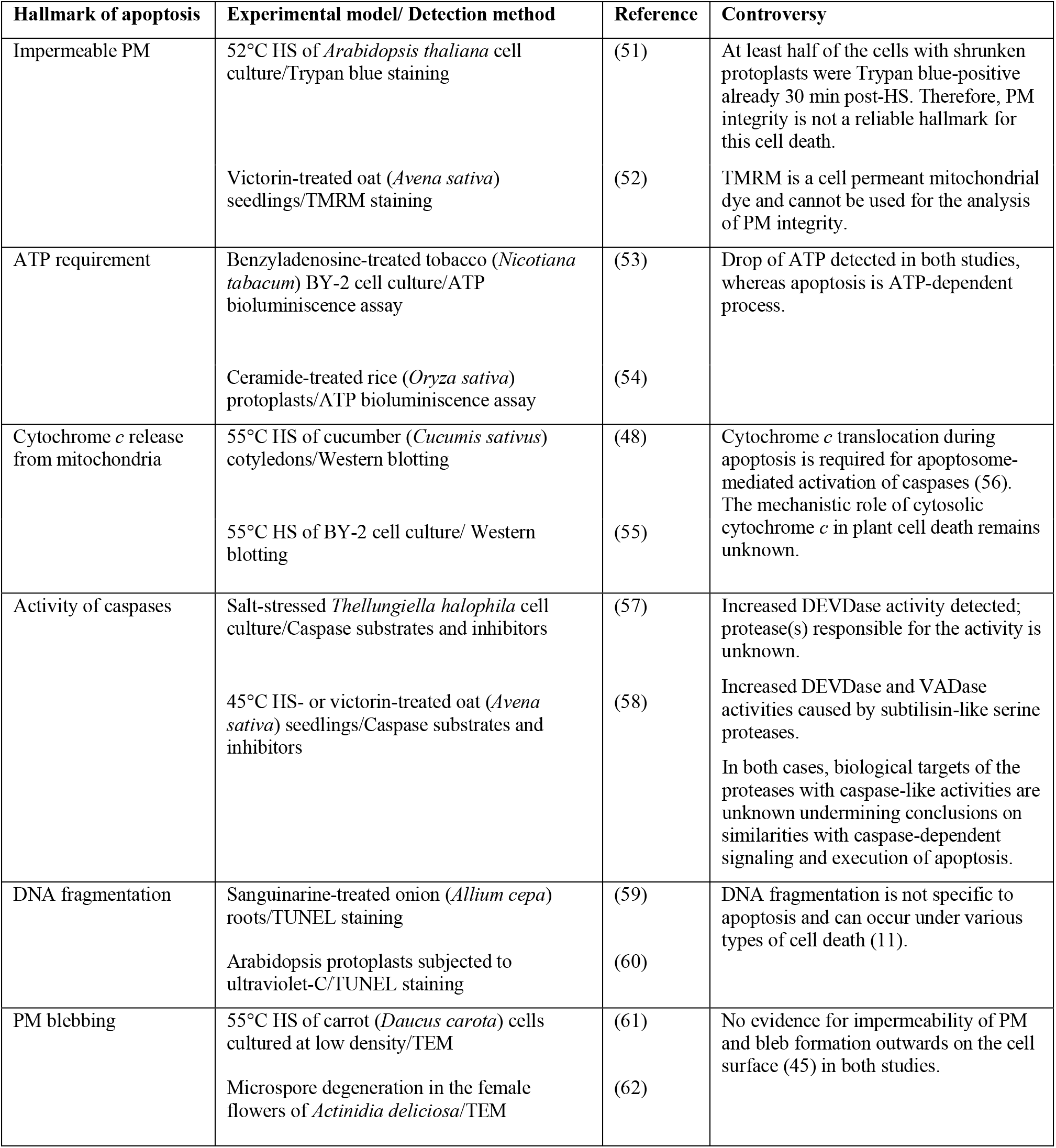
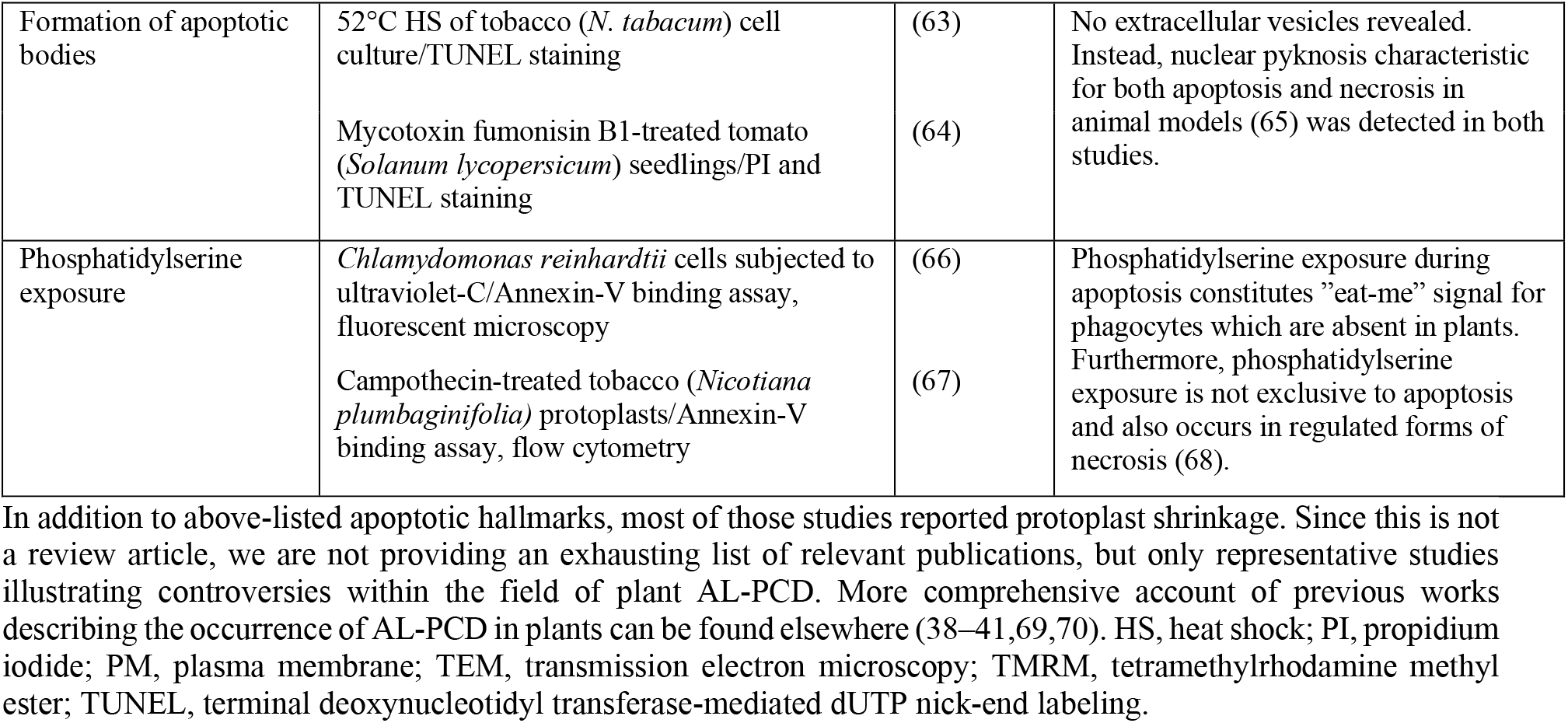
A brief account of major hallmarks of animal apoptosis analyzed in plant models for AL-PCD.

Plant AL-PCD is commonly induced by an acute heat stress, e.g. a pulse heat shock (HS) at 40-55°C (**Table 1**; (49,71–74)). However, such temperatures are on the edge of the physiologically relevant range and could likely cause acute damage resulting in accidental necrotic cell death, rather than trigger a programmed response. Unlike apoptosis, accidental necrotic cell death is associated with a drop of cellular ATP content, occurs independently of caspases, displays early PM permeabilization leading to the leakage of the cellular content into the extracellular environment and is considered to be generally unprogrammed process under conditions when RCD fails (75,76).

To reconcile the reported features of plant AL-PCD with the confirmed lack of the key components of the apoptotic machinery and the general ineptness of the apoptotic cell death strategy in the context of a plant organism, we systematically analysed cell death hallmarks occurring under conditions reported to trigger AL-PCD (40-55°C HS) and necrosis (85°C HS) (61,70). We took special care to include appropriate controls for potential technical pitfalls and performed most of the experiments using highly reliable Bright Yellow-2 (BY-2) tobacco cell culture, the so-called “plant HeLa” model system (77) often implemented for plant RCD studies (**Table 1**; (49,78)). We show that while protoplast shrinkage during 55°C HS-induced cell death superficially resembles AVD, it is independent of ATP or Ca^2+^ and coincides with instant and irreversible loss of PM integrity. This cell death lacks other key apoptotic features, such as PM blebbing, nuclear segmentation, and formation of apoptotic bodies. The side-by-side comparison of cell death triggered by 55°C and 85°C revealed remarkable similarities to necrosis in both cases. Importantly, the above-described hallmarks of accidental necrosis would also match features of regulated necrotic cell death (79). However, this study did not aim at determining whether necrosis induced by HS is accidental or regulated process. Furthermore, we demonstrate that the previously reported striking morphological difference between cell deaths triggered by 55°C and 85°C, i.e. the lack of protoplast shrinkage in cells exposed to 85°C, is merely a consequence of the fixative effect of the high temperature on cellular content. We conclude that cell death in the model system typically used to study AL-PCD is indistinguishable from necrosis.

## Results and discussion

### HS-induced plant cell death is morphologically distinct from apoptosis

One of the earliest hallmarks of apoptosis is AVD (80,81), followed by blebbing of the PM, nuclear segmentation and finally fragmentation of the cell into apoptotic bodies (17). Since the ultimate goal of such cell dismantling is removal of the dying cell by phagocytes, the PM of the apoptotic cell should remain intact throughout the whole process, preventing spillage of the dying cell content. It is fairly obvious that formation of apoptotic bodies during plant cell death would serve no purpose, as plants lack phagocytes and a dead plant cell remains to be incapsulated within a rigid cell wall (25,82). However, death of plant cells caused by abiotic or biotic stress is accompanied by rupture of PM and/or vacuolar membrane, tonoplast, causing severe damage to the endomembrane systems and irreversible detachment of the PM from cell wall which is often referred to as the protoplast shrinkage (11,83). This phenomenon has been repeatedly confused with AVD, PM blebbing and formation of apoptotic bodies, and thus exploited to support existence of AL-PCD in plants (38,39).

To examine AVD, PM blebbing, nuclear segmentation and cellular fragmentation in plants, we analysed morphology of tobacco BY-2 cells under HS conditions that were previously described to induce AL-PCD (48,51). Cell culture was stained with a noncell-permeable Sytox Orange (SO) nucleic acid dye to visualize cells with compromised PM integrity and with the styryl dye FM4-64 to visualize cell membranes and then subjected to a pulse HS at 55°C (**Additional file 1: Video S1**).

Only SO-positive cells exhibited protoplast shrinkage, strongly indicating correlation between cell volume decrease and PM permeabilization (**Fig. 1a)**. Notably, the treated cells became SO-positive at the earliest checked time point, i.e. within 10 min of the HS indicating rapid PM permeabilization. One prominent feature of SO-positive cells was formation of vesicle-like structures at the inner side of the PM (**Fig. 1a, b)**. Such morphology is inconsistent with AVD and blebbing, which require intact PM; furthermore, apoptotic PM blebs form outwards on the cell surface (45).

**Figure 1.**
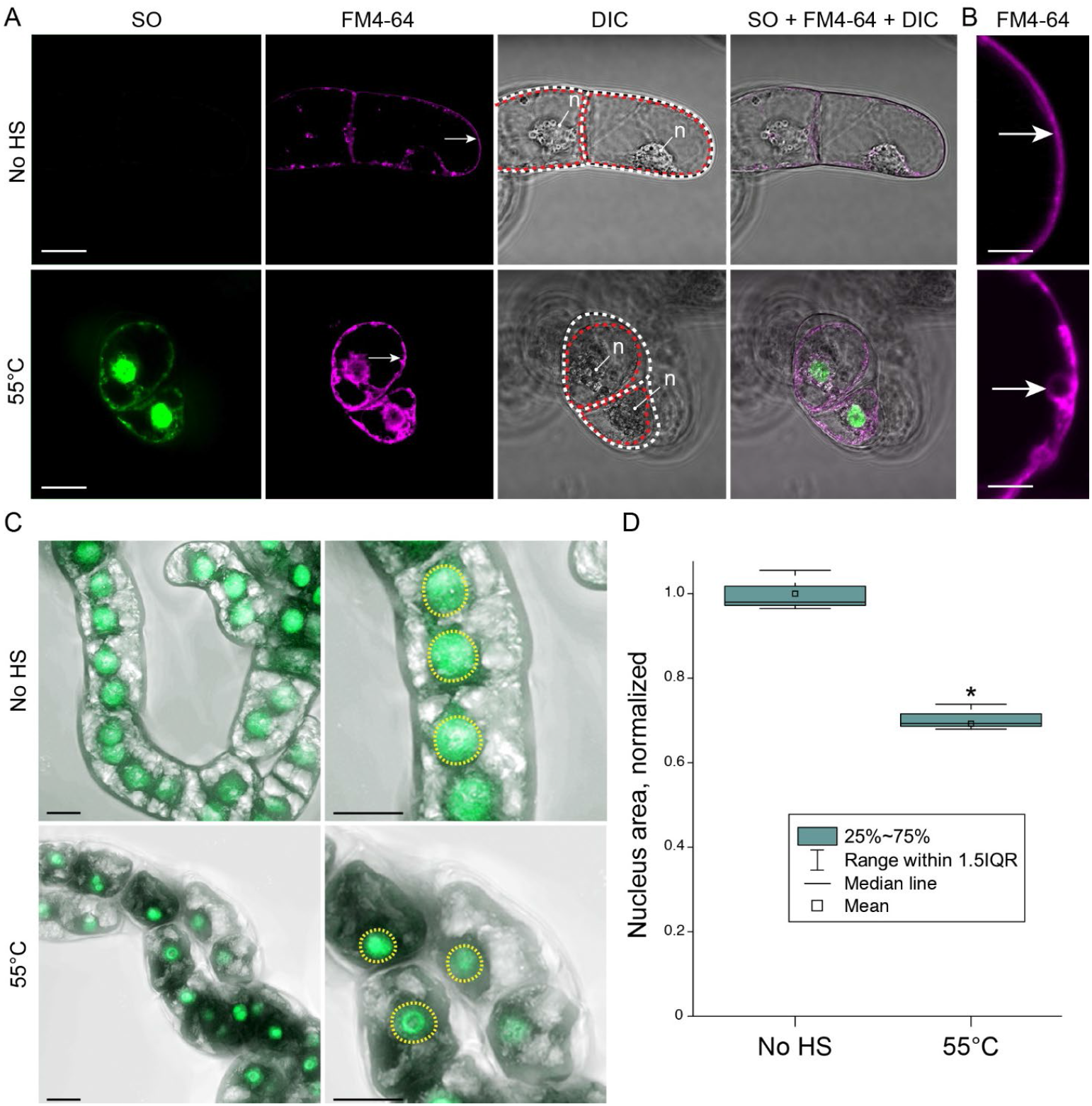
Gross morphological changes in HS-treated plant cells do not match hallmarks of apoptosis. **a** FM4-64 and SO dyes were used to visualize all cells and cells with permeabilized PM, respectively. BY-2 cells were imaged under control conditions (No HS) or after a 10-min HS at 55°C and were stained according to protocols i and ii, respectively (see Methods, *Sytox Orange and FM4-64 staining*). Scanning was performed within 1 h post-HS. The arrows indicate PM; note the PM (red dotted line) being tightly pressed to the cell wall (white dotted line) under control conditions and detached under stress conditions. **b** Higher magnification of the areas indicated with arrows in **a**. **c** BY-2 cells expressing green fluorescent protein (GFP) fused to nuclear localization signal (NLS) were subjected to the same treatments as in **a**. Images represent maximum intensity projections of z-stack scans acquired 6 h post-HS. **d** Quantification of nuclear area in samples shown in **c**. Data from three independent experiments, with ≥ 142 cells counted per treatment. Student’s t-test, *p < 0.005. DIC, differential interference contrast microscopy. n, nucleus. IQR, interquartile range. Scale bars, 20 μm (**a, c**) or 5 μm (**b**).

The PM and shrunken protoplast were additionally imaged in the time interval from 15 min to 72 h after the pulse HS (**Additional file 2: Fig. S1**). Nevertheless, we failed to detect PM blebbing or protoplast fragmentation into discrete bodies at any time point. Furthermore, although we did observe moderate nuclear condensation upon HS, it was not followed by segmentation of the nucleus (**Fig. 1c, d**). In summary, the gross morphological changes of cells undergoing HS-induced cell death do not resemble apoptosis.

### PM integrity is irreversibly compromised during or shortly after pulse HS

Intact PM is a pivotal hallmark of apoptosis (10). However, SO staining described above suggested permeabilization of PM in most BY-2 cells already within 10 min of the HS. To investigate dynamics of the PM permeabilization, we analysed cellular content leakage after two types of HS, at 55°C or 85°C which were reported to induce AL-PCD or necrosis, respectively (61,70). The leakage of cellular content was assessed using a fluorescein diacetate (FDA)-based fluorochromatic assay (84). In brief, the non-fluorescent FDA molecules can passively diffuse into the living cells where their acetate groups are cleaved off by esterases. The resulting fluorescein molecules have poor membrane permeability and are retained in cells with an intact PM, but are released into extracellular space upon PM permeabilization.

We imaged BY-2 cells loaded with FDA prior to the pulse HS at 55°C or 85°C (**Fig. 2a**). Both types of HS caused rapid (within 10 min) leakage of the dye into the extracellular space (**Fig. 2a**). We measured the amount of fluorescein accumulated in the extracellular space immediately after the HS and found that the rate of cellular content leakage in HS-treated cells was comparable to that occurring after severe disruption of plant cells caused by freeze-thaw in liquid nitrogen (**Fig. 2b**).

**Figure 2.**
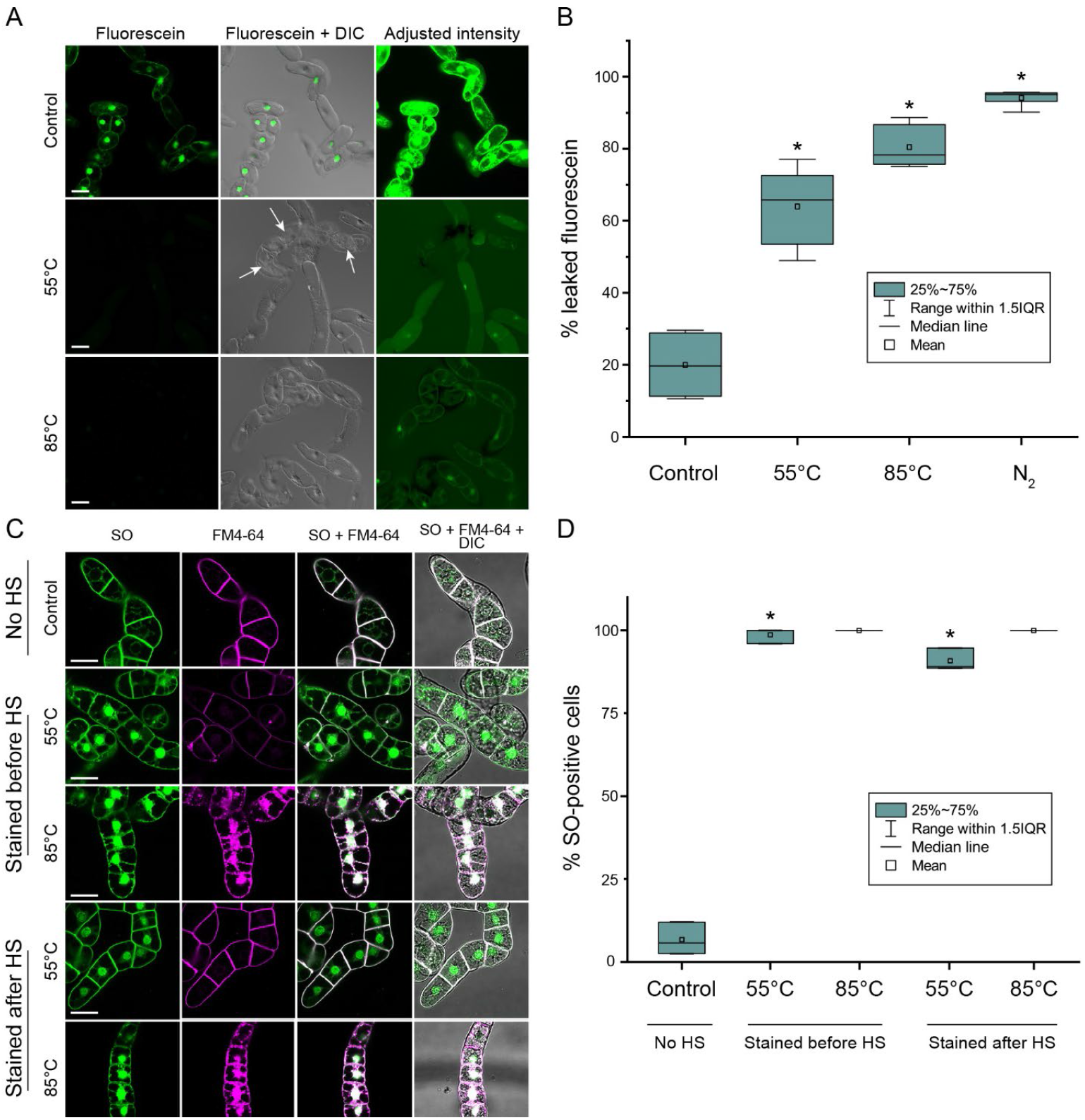
Both 55°C and 85°C HS cause instant and irreversible PM permeabilization. **a** Green fluorescence in the BY-2 cells loaded with FDA. Fluorescein leaks into extracellular space within 10 min after HS at either 55°C or 85°C. Arrows indicate shrunken protoplasts in 55°C HS-treated cells. **b** Similar fraction of fluorescein leaks out of cells after 10 min of 55°C, 85°C, or liquid nitrogen (N_2_) treatment. **c** BY-2 cells stained with SO and FM4-64 to assess whether disruption of the PM integrity after pulse HS is transient or irreversible. Staining was performed for no HS, before HS or after HS treatments according to protocols i, ii and iii, respectively (see Methods, *Sytox Orange and FM4-64 staining*). The cultures were imaged within 1 h following HS. **d** Frequency of the SO-positive cells in the cultures shown in **c**. DIC, differential interference contrast microscopy. IQR, interquartile range. Scale bars in **a** and **c**, 50 μm. **b**, representative data from one out of three independent experiments. **d**, data from three independent experiments, with ≥ 115 cells counted per treatment. **b** and **d**, one-way ANOVA with Dunnet’s test; *p<0.005.

To determine whether PM permeabilization was transient or permanent, we added SO and FM4-64 stains to the cell cultures either before or 30 min after HS. We speculated that if PM permeabilization upon HS was transient, cultures stained after HS would show a significantly lower frequency of SO staining as compared to cells stained before HS. However, SO staining before and after 55°C or 85°C HS showed no differences in the proportion of SO-positive cells (**Fig. 2c, d**), indicating that both treatments caused irreversible rapid permeabilization of PM typical for necrosis (10,11,85).

### The protoplast shrinkage during HS-induced cell death is ATP- and Ca^2+^-independent

Dismantling of the apoptotic cells is ATP-dependent (42,86). Yet, loss of the PM integrity would lead to rapid depletion of intracellular ATP, rendering all energy-dependent processes defunct. To examine whether the HS-induced cell death requires ATP, we first imaged mitochondria after 55°C and 85°C HS. Already at the earliest checked time points 4-10 min after HS, under both temperatures, mitochondrial dye MitoTracker localized to aberrant structures similar to those observed after treatment with mitochondrial uncoupler and ATP synthesis inhibitor, protonophore carbonyl cyanide m‐chlorophenylhydrazone (CCCP) (87), indicating disruption of mitochondrial membrane potential (MMP; **Fig. 3a**). Furthermore, intracellular ATP content dropped dramatically after both HS treatments (**Fig. 3b**), most probably due to dissipation of the MMP and leakage of cytoplasmic content through the permeabilized PM.

**Figure 3.**
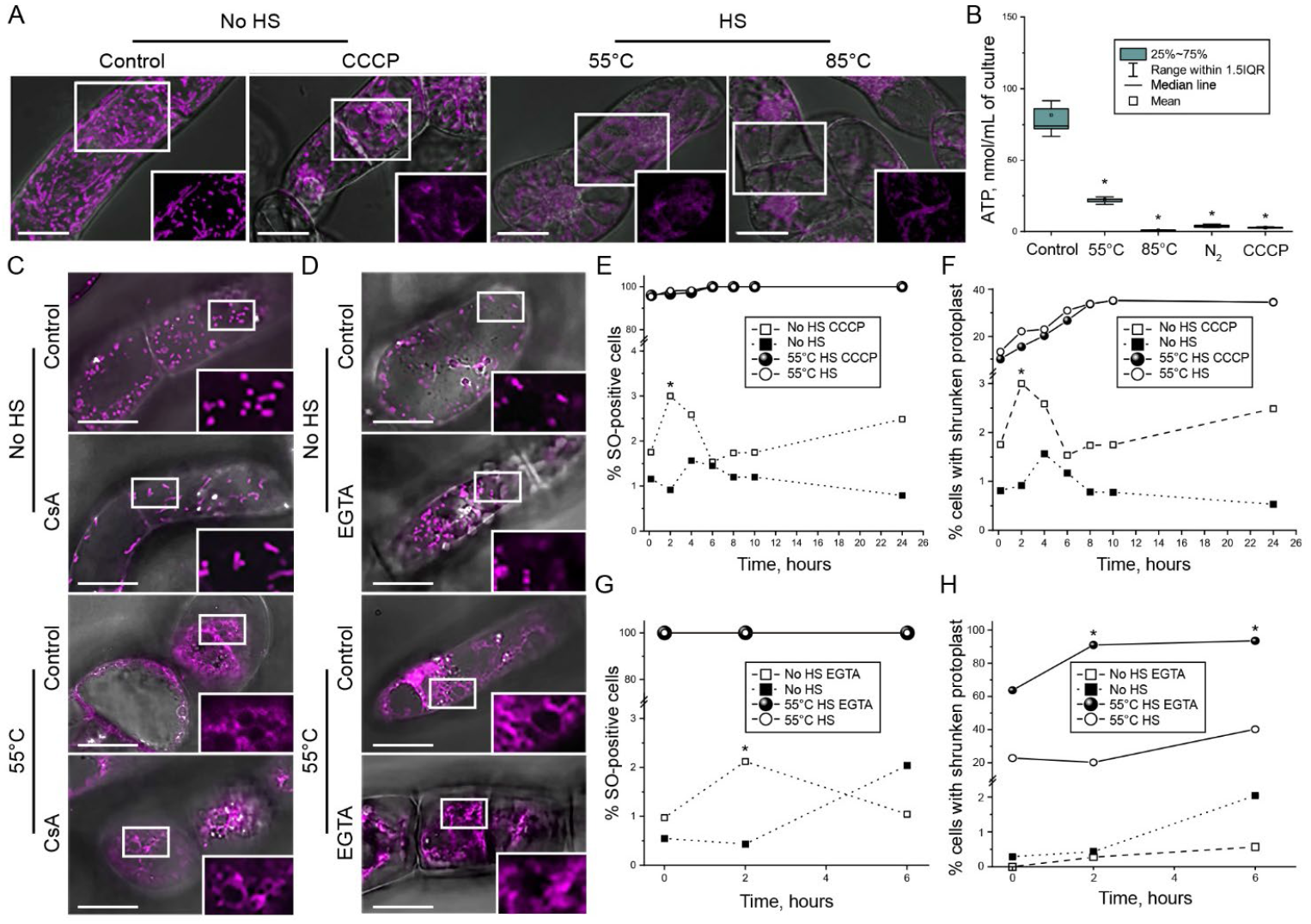
Protoplast shrinkage at 55°C HS is an ATP- and Ca^2+^-independent process. **a** Mitochondria in BY-2 cells stained with MitoTracker Red and imaged 10 min after 55°C or 85°C HS, or after treatment with 48 μM CCCP under normal temperature. Severely damaged mitochondria were observed upon all three treatments. **b** Loss of intracellular ATP content upon HS. Snap freeze-thaw treatment in liquid nitrogen (N2) and CCCP treatment were used as positive controls for completely disrupted and uncoupled mitochondria, respectively. The experiment was repeated twice, each time using four biological replicates per treatment. **c** MitoTracker Red staining of BY-2 cells exposed to 55°C in the presence or absence of 15 μM Cyclosporin A (CsA) reveals that inhibition of MPTP opening does not rescue mitochondria from severe damage and loss of MMP caused by HS. **d** MitoTracker Red localization in the cells pre-treated with 10 mM EGTA prior to the HS reveals that chelation of extracellular Ca^2+^ does not rescue mitochondrial phenotype. **e, f** Dynamics of cell death (% SO-positive cells; **e**) and protoplast shrinkage **(f)** in cells with normal and uncoupled (48 μM CCCP treatment) mitochondria. **g, h** Pre-treatment with 10 mM EGTA before HS does not affect dynamics of cell death (% SO-positive cells; **g**) and protoplast shrinkage **(h)**. Experiments shown in **e-h** were repeated three times, with ≥ 184 cells per treatment and time point. Each microscopy experiment was performed at least twice. Staining for no HS and 55°C treatments were performed according to protocols i and ii, respectively (see Methods, *Sytox Orange and FM4-64 staining*). Scale bars, 20 μm (**a**) or 50 μm (**c, d**). IQR, interquartile range. **b**, **e**-**h**, one-way ANOVA with Dunnet’s test; *p<0.05.

In addition to ATP depletion, PM permeabilization would also cause entry of Ca^2+^ into the cells, potentially followed by its accumulation in mitochondria. MMP-driven accumulation of Ca^2+^ can trigger mitochondrial permeability transition (MPT) due to the opening of a nonspecific pore (mitochondrial permeability transition pore, MPTP) (88), which will cause arrest of ATP synthesis and production of reactive oxygen species ultimately resulting in necrotic cell death (89). Although HS-induced plant cell death was previously suggested to be a Ca^2+^-dependent process (51), those experiments lacked controls for mitochondrial phenotype, respiration, or ATP production.

To test whether MPT plays a role in the observed mitochondrial phenotype, experiments were performed in the presence of cyclosporin A (CsA), an inhibitor of MPTP opening, or the Ca^2+^ chelator ethylene glycol-bis(2-aminoethylether)-N,N,N’,N’-tetraacetic acid (EGTA). Neither CsA nor EGTA could alleviate the mitochondrial phenotype in cells subjected to the HS (**Fig. 3c, d**), indicating that mitochondrial malfunction was caused by the direct loss of the mitochondrial membrane integrity during the HS, independently on PM permeabilization.

Next, we examined the importance of intracellular ATP for the AVD-like protoplast shrinkage. We found that pre-treatment of the cell cultures with CCCP prior to HS had no effect on the protoplast shrinkage (**Additional file 2: Fig. S2**), demonstrating that the protoplast shrinkage does not require ATP. Time-resolved quantitative analysis of cell death (frequency of SO-positive cells) and protoplast shrinkage upon HS of cells with normal or uncoupled mitochondria revealed that both parameters were independent of the mitochondrial bioenergetic function (**Fig. 3e, f**). Pre-treatment with CsA or EGTA prior to HS did not alleviate the cell death rate either **(Fig. 3g, h**; **Additional file 2: Fig. S2**).

The discrepancies between our observations and the previous studies could be caused by technical issues, primarily precision of the temperature measurement during HS. To examine this possibility, we compared frequency and phenotype of cell death after treatment at 40, 45, 50 and 55°C. Although cell viability was inversely proportional to the temperature, the morphology of dead cells in all cases was identical to that observed at 55°C (**Fig. 4**). Pre-treatment of cells with CCCP confirmed that at any of the checked temperatures the rate of cell death did not depend on mitochondrial activity (**Fig. 4**).

**Figure 4.**
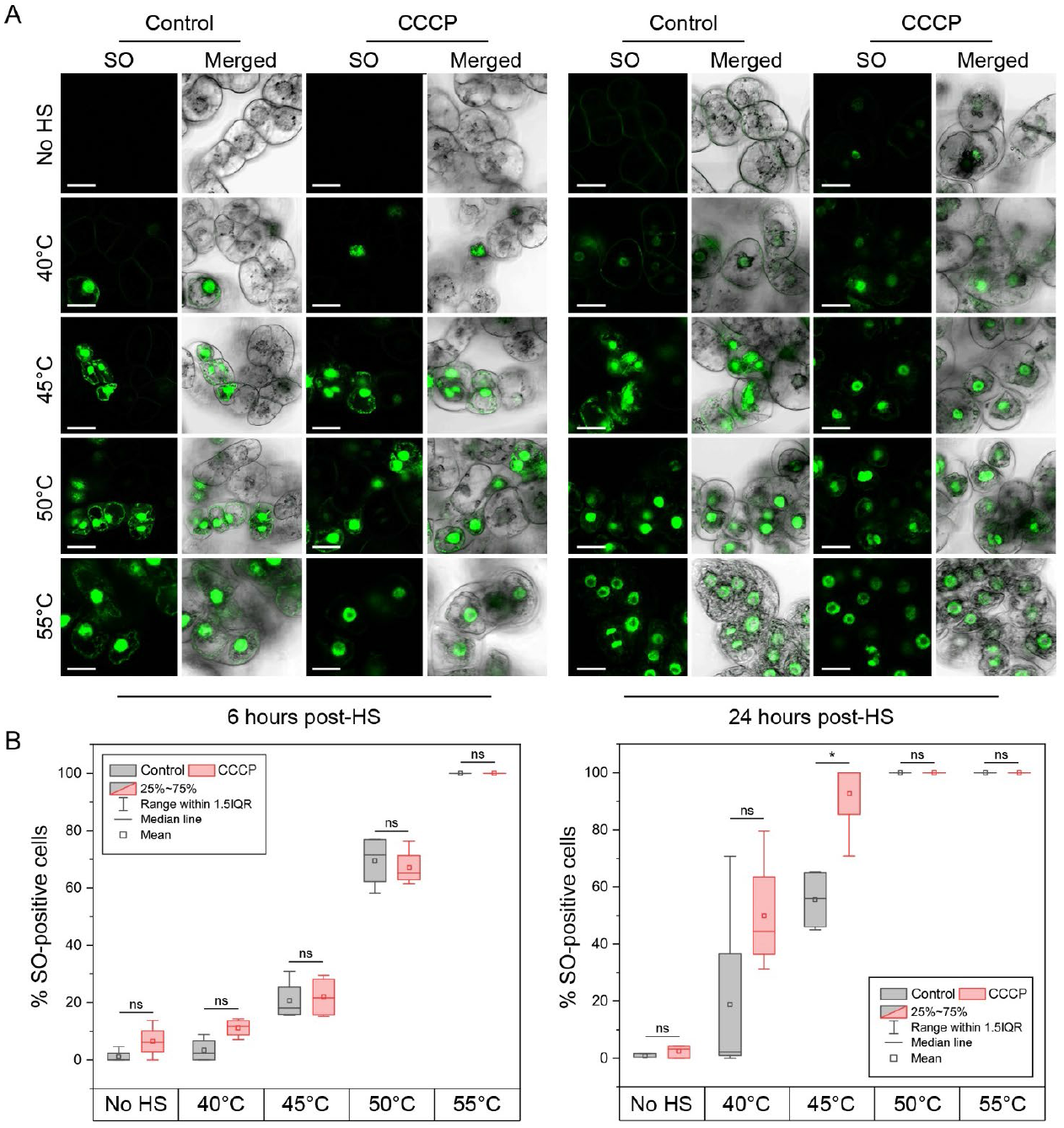
HS at the temperature range 40-55 ° C induces ATP- independent cell death. **a** SO staining of BY-2 cells heat-shocked for 10 min at 40, 45, 50, or 55°C and imaged after 6 or 24 h. **b** Quantification of cell death (% SO-positive cells) in the samples showed that pre-treatment with CCCP provided no protection against cell death and protoplast shrinkage at any of the tested HS temperatures. On the contrary, after prolonged exposure (24 h), CCCP appeared to decrease HS tolerance of plant cells. Staining was performed for no HS and HS treatments according to protocols i and ii, respectively (see Methods, *Sytox Orange and FM4-64 staining*). DIC, differential interference contrast microscopy. IQR, interquartile range. Experiments were repeated three times, with ≥ 134 cells per treatment and time point. The data was subjected to one-way ANOVA with Bonferroni correction. *p<0.05, ns, non-significant. Scale bars, 20 μm.

A recent study (90) proposed that HS at 55°C caused ferroptosis in *A. thaliana* root hair cells. Although mitochondrial dysfunction is known to suppress ferroptosis in animal cells (91) and, as shown above, HS-induced plant cell death is mitochondria-independent, we still examined whether this cell death could be classified as a ferroptosis. For this, BY-2 cells were treated with a ferroptosis inhibitor, Ferrostatin-1 (Fer-1), prior to HS at 55°C and cell death rate was measured during 24 h after the stress. Treatment with Fer-1 did not alleviate the cell death (**Fig. 5a, b;** data is shown for the first 12 h after HS). Furthermore, two independent experiments replicating conditions that were reported to induce ferroptosis in *Arabidopsis* root hair cells (90) did not confirm such type of cell death (**Fig. 5c)**. Taken together, our results reject the notion that HS-induced cell death is a programmed process. On the contrary, they demonstrate that HS triggers rapid destruction of cellular components and passive decay of plant cells resembling accidental necrosis.

**Figure 5.**
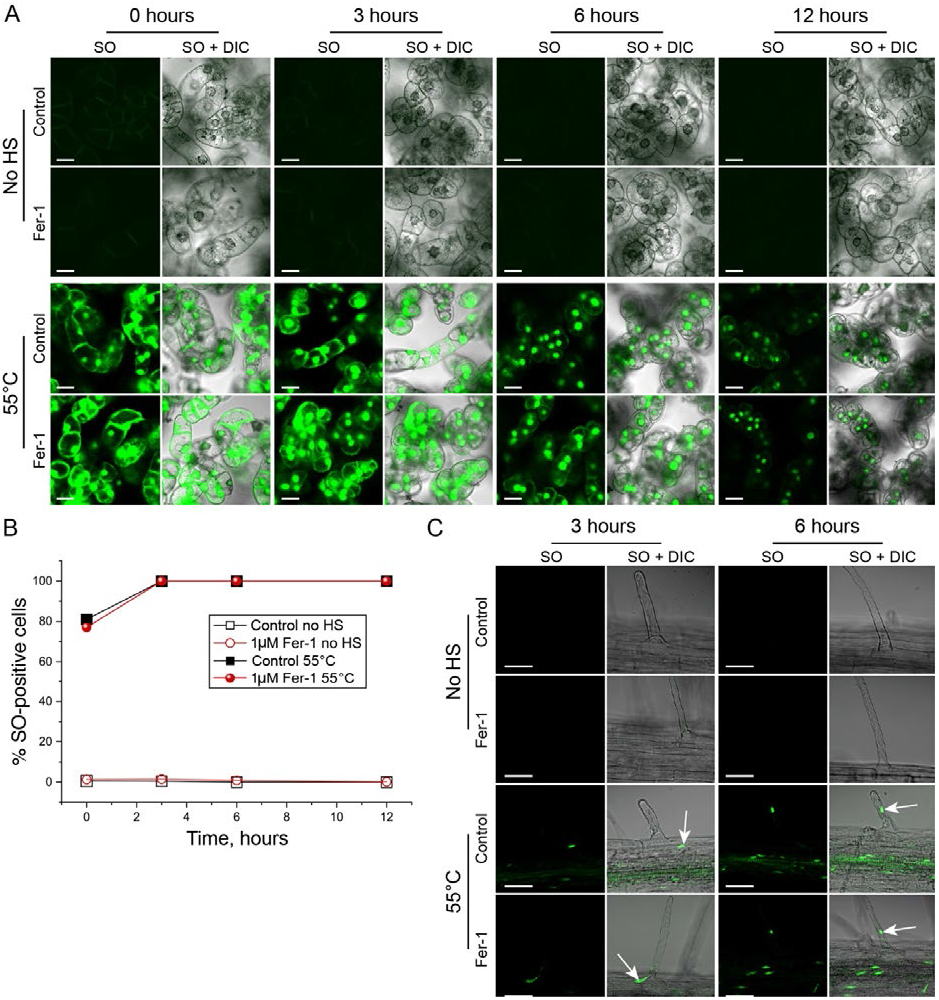
HS-induced cell death response is not ferroptosis. **a** SO staining of BY-2 cell cultures demonstrates that pre-treatment with 1 μM Fer-1 does not affect HS-induced cell death. Staining for no HS and 55°C treatments was performed according to protocols i and iii, respectively (see Methods, *Sytox Orange and FM4-64 staining*). **b** Quantification of cell death frequency in the samples illustrated in **a**. The chart shows representative results of three independent experiments, each including ≥280 cells per treatment and time point. **c** Pre-treatment with Fer-1 does not alleviate the HS-induced death of *Arabidopsis thaliana* root hair cells. Arrows indicate SO-positive nuclei. Three independent experiments demonstrated the same results. No quantification was performed in these experiments, since all cells were SO-positive. DIC, differential interference contrast. Scale bars, 20 μm

### Necrotic deaths caused by 55°C or 85°C display different cell morphologies due to a fixating effect of higher temperature

The lack of protoplast shrinkage during cell death induced by 85°C was used to classify it as necrosis (61,70). However, both 55°C and 85°C HS trigger instant and irreversible permeabilization of the PM, MMP dissipation, and drop in intracellular ATP content, which are hallmarks of necrosis (85). Plausibly, morphological differences between necrotic cell deaths triggered by 55°C and 85°C could be explained by rapid protein denaturation occurring at 85°C that would crosslink cellular components. Such “fixation” would prevent protoplast shrinkage. Consistent with this suggestion, high-temperature cell fixation protocols have been used as an alternative to chemical fixation (92).

To test whether 85°C HS acts as a fixative, we induced protoplast retraction from the cell wall by exposing stressed cell cultures to hypertonic conditions at 500 mM D-mannitol (**Fig. 6a**). The high osmotic pressure of the medium would induce dehydration and protoplast shrinkage in the non-fixed cells. As expected, the living cells treated with D-mannitol underwent typical plasmolysis manifested by reversible protoplast detachment from the cell wall. Cells exposed to 55°C displayed irreversible protoplast shrinkage phenotype both with and without D-mannitol treatment. However, although cells treated at 85°C HS exhibited visible signs of dehydration in the hypertonic solution, protoplasts of virtually all cells remained attached to the cell wall. Quantitative analysis revealed no significant changes in the protoplast area of cells treated at 85°C HS under normal or high osmotic pressure (**Fig. 6b**). These data provide compelling evidence that the fixing effect of 85°C HS prevents protoplast shrinkage. Thus, distinct phenotypes of cell deaths induced by HS at 55°C and 85°C do not reflect differences in the cell-death execution mechanism.

**Figure 6.**
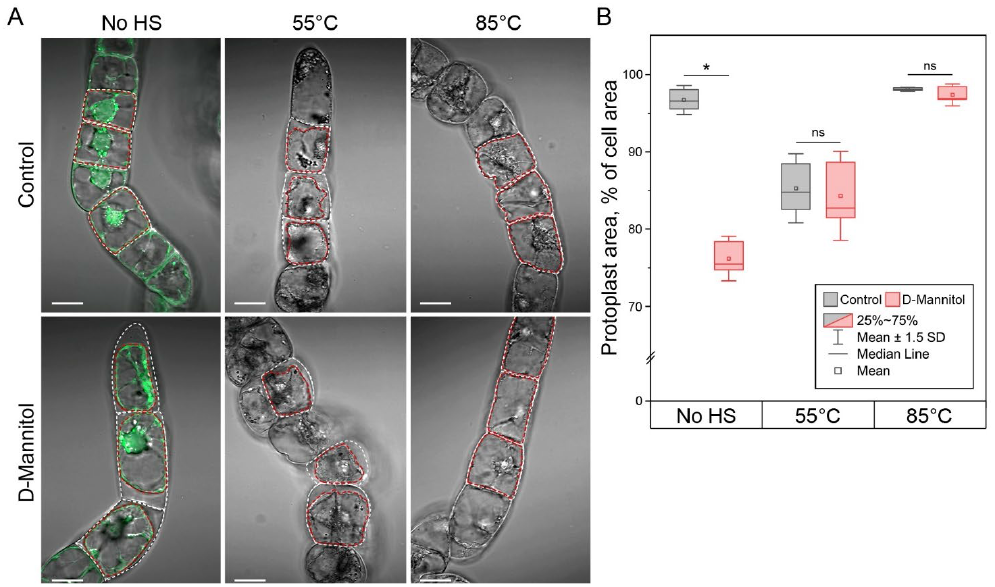
85°C HS prevents protoplast shrinkage in necrotic cells by fixing cellular content. **a** Cells stained with FDA were exposed to 55°C or 85°C and mounted in the normal growth medium or hypertonic medium supplemented with 500 mM D-mannitol. As expected, hypertonic medium induced plasmolysis in the non-stressed cells (No HS) and had no effect on protoplasts shrunken after 55°C HS. Importantly, high osmotic pressure failed to induce protoplast shrinkage in cells exposed to 85°C HS, thus confirming that cells were fixed by the high-temperature treatment. **b** The extent of protoplast shrinkage upon HS followed by increased osmotic pressure was quantified as the percentage of the cell area (outlined by the white dotted line in **a**) occupied by the protoplast area (outlined by the red dotted line in **a**). Student’s t-test, n = 3 replicates (each replicate representing ≥ 34 cells), *p<0.0001, ns, non-significant. IQR, interquartile range. Scale bars, 20 μm.

## Conclusions

This study draws attention to the existing disagreements in plant cell-death research community about existence of apoptosis in plants and emphasizes the need for in-depth investigation of plant-specific RCD features, which will hopefully boost our understanding of the evolution of eukaryotic cell death machinery. Research on plant cell death greatly benefits from the wealth of knowledge obtained on animal RCD, but also frequently suffers from the lingering superficial comparisons between both systems as exemplified in **Table 1**. Extrapolating detailed knowledge about animal RCD onto plant processes poses a tempting shortcut to a seemingly meaningful interpretation of fragmented data available for plant model systems. Albeit such comparisons can be helpful to promote a new line of investigation, caution should be exercised to ensure that they provide a functional insight and do not lead to a logical dead-end, as it happened with the reports on formation of apoptotic bodies in plants.

An interesting illustrative example of such cautious comparison are the recent works revealing similarities in the complex formation by Nod-like receptor (NLR) proteins involved in plant and animal innate immune responses and cell death regulation (93–95). These studies showed that resistosome complexes formed by plant NLRs structurally resemble the inflammasome, complex of animal NLRs assembled during immune response and even apoptosome complex formed by the Apaf-1 protein during apoptosis (56,96,97). These complexes share a multimeric wheel-like shape; furthermore, both plant resistosomes and animal inflammasomes are proposed to trigger cell death mediated by formation of pores in the PM. However, this exciting resemblance is at the same time juxtaposed by a fundamental difference in the mechanisms of action for these complexes in plant and animal cell deaths. While plant resistosomes are proposed to directly form pores in the PM (97), the animal inflammosome and apopotosome complexes evolved to work as platforms for activation of caspase cascades in the cytoplasm (56,96).

Results of our current work schematically summarized in **Fig. 7** indicate that morphological and biochemical characteristics of plant HS-induced cell death termed in the literature AL-PCD *de facto* fit the definition of necrosis. Here we exampled a detailed analysis of only one typical model for plant AL-PCD. Although, we cannot rule out that other stimuli could trigger RCD with apoptotic features, this scenario is unlikely due to the lack of the core apoptotic machinery in plants and general pointlessness of apoptosis relying on clean up by mobile cells in the context of plant biology. The set of assays described here can be easily implemented for other plant model systems used for the investigation of cell death with apoptosis-like features (**Table 1**; (59,74,98–108)) and will hopefully provide insights for still existing uncertainties.

**Figure 7.**
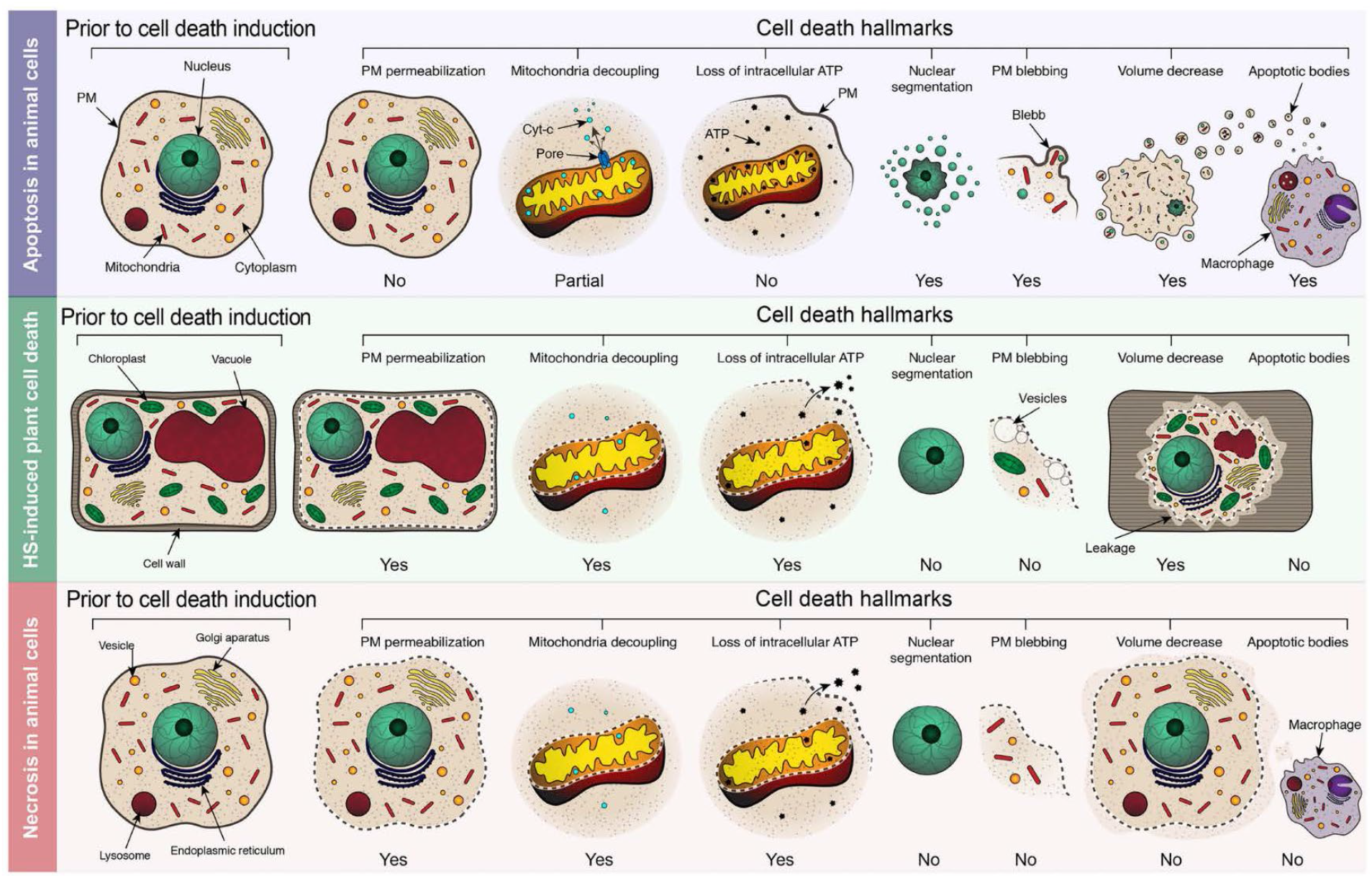
HS-induced plant cell death is necrotic rather than apoptotic process. Plasma membrane (PM) integrity is maintained during apoptosis to prevent spillage of toxic dying cell contents. However, plant cell exposed to HS and undergoing the so-called AL-PCD exhibit early irreversible permeabilization of PM, a feature typical for necrotic cell death. Apoptosis is associated with step-wise massive re-organization of the cellular morphology (nuclear segmentation, PM blebbing, fragmentation into apoptotic bodies) and thus is an energy consuming process that relies on controlled release of cytochrome *c*, functional mitochondria and intracellular ATP. On the contrary, necrotic death is a passive lysis of the cellular contents associated with low intracellular ATP and decoupled mitochondria. Similarly to animal cells undergoing necrosis, plant cells subjected to HS have a low intracellular ATP content and uncoupled mitochondria. The PM of apoptotic cell forms outward protrusions, called blebs, that will eventually undergo scission and become apoptotic bodies. Vesicles that are observed in the proximity of plant PM during HS-induced cell death are on the inner side of the PM and thus cannot be considered as blebs. While apoptotic cells condense and disassemble into apoptotic bodies that are engulfed by phagocytes, a necrotic cell typically undergoes swelling that is followed by a decrease of the cell volume and leakage of the dead cell contents. Presence of the rigid cell wall most probably does not permit necrosis-like swelling of plant cells during HS-induced cell death, while rupture of the vacuole coinciding with permeabilization of the PM would lead to rapid leakage of the cellular contents and thus decrease in volume. Such shrinkage has very little in common with AVD, as it is a passive process not associated with the formation of apoptotic bodies.

## Methods

### Plant material and growth conditions

BY-2 cell cultures (77) were grown at 25°C (unless stated otherwise), 120 rpm, in the darkness. Cells were subcultured every 7 days to the standard Murashige and Skoog (MS) medium supplemented with vitamins (M0222, Duchefa), 1.875 mM KH_2_PO_4_, 0.2 mg/L 2,4-D, and 3% (w/v) sucrose; pH 5.6.

For growing *Arabidopsis thaliana* seedlings, Col-0 seeds were sterilized for 30 min using a 2.7 g/L sodium hypochlorite solution (Klorin, Colgate Palmolive) with 0.05% (v/v) Tween 20. The seeds were then rinsed three times with filter-sterilized Milli-Q water prior to being plated on solidified medium. The MS medium was brought to pH 5.8 with 1M KOH prior to autoclaving and contained half-strength MS salts and vitamins (M0222, Duchefa), 1% (w/v) sucrose, 10 mM MES and 0.8% (w/v) plant agar (P1001, Duchefa). The plates were incubated vertically with cycles of 16 h of 150 μE m^−2^ s^−1^ light at 22°C/8 h of dark at 20°C.

### Fluorescein leakage assay (fluorochromatic assay)

Cells were stained with 4 μg/mL FDA in the culture medium for 10 min at room temperature. Half a mL aliquots of the stained cell culture were then transferred into 2 mL Eppendorf tubes using a cut 1-mL tip to avoid mechanical damage of the cells. Each tube was treated as a biological replicate. At least three replicates were used for each condition in each experiment.

The tubes were exposed to four conditions: (i) left on the bench at room temperature; incubated for 10 min at (ii) 55°C or (iii) 85°C in a preheated thermoblock (wells were filled with water for better thermoconductivity); (iv) snap frozen in liquid nitrogen. After treatments, cultures were let to cool down or thaw for 5 min at room temperature and then either mounted on a sample glass to be imaged using confocal laser scanning microscopy (CLSM) or processed further for quantitative assay.

For quantitative fluorescein leakage assay cells were pelleted on nylon meshes with 50 μm pores. Supernatants were collected entirely with a pipette, 150 μL of each supernatant were transferred to the empty wells of a 96-well flat bottom plate (Sarstedt). Cells on the mesh were then washed off the meshes into wells of the same plate using 150 μL of fresh BY-2 media. Fluorescence was measured in the FLUOstar Omega Microplate Reader (BMG LABTECH) using 485 nm excitation/520 nm emission filters (gain 800, 20 flashes/well). Importantly, the measurement had to be performed within 20 min after treatment to avoid passive diffusion of fluorescein into extracellular space in the control samples. Leaked fluorescein intensity was calculated for each sample as a percentage of total fluorescence intensity detected in the cells and supernatant. A one-way ANOVA with a Dunnet’s test was performed using JMP 10, and the graph was built in Origin2019.

### Sytox Orange and FM4-64 staining

Aliquots of 4-5-day old BY-2 cell culture were transferred to 1.5 mL Eppendorf tubes using a cut 1 mL tip. The final concentrations of Sytox Orange (SO) and FM4-64 in cell culture were 1 μM and 0.5 μM, respectively. Depending on experimental design, four different staining protocols were applied to the cell cultures: (i) As a control, stained cell culture was incubated on the lab bench for 30 min. (ii) Stained cell culture was incubated in a preheated thermoblock for a 10-min HS. (iii) Cell culture was subjected to a HS for 10 min, and stained after additional 30 min at room temperature. (iv) For multi-day (72 h) experiments, cell culture was subjected to a HS for 10 min, and stains were applied 10 min before observation. Cells were mounted on a microscope slide and imaged by CLSM after staining. More than one hundred cells were counted for each treatment to ensure robustness of the results. Note that there is some degree of crosstalk in the fluorescence emission profiles of the SO and FM4-64 stains, which was most prominent following 85°C HS; however this crosstalk had no influence on the results of our experiments.

### Nuclear phenotyping

A transgenic BY-2 line expressing GFP fused to NLS under a double 35S promoter (2×35::Rluc-sGFP-NLS) was used to evaluate nuclear phenotype following HS. The transgenic culture was generated according to (109). Treatments included: (i) No HS, (ii) 10-min pulse HS at 55°C (iii) 10-min pulse HS at 55°C with SO staining as stated above, which was used to validate results following HS (data not shown). Six hours following HS, z-stack acquisitions were captured with Zeiss LSM 800 as described in the *Confocal microscopy*. Quantification was performed using maximum intensity projections. The areas of nuclei were approximated using threshold segmentation of images with Fiji software (ImageJ version 1.52r). Three independent experiments were carried out.

### MitoTracker Red Staining

BY-2 cells were stained with 100 μM MitoTracker Red (ThermoFisher, M7512) for 10 min at room temperature. Cells were mounted on a microscope slide and imaged using Zeiss LSM 780 as described in the *Confocal microscopy*.

### EGTA and cyclosporin A treatments

Half a mL of a 4-5-day old BY-2 cell culture were transferred to 1.5 mL Eppendorf tubes. The cells were stained with MitoTracker Red (100 μM) and FDA (2 μg/mL) 10 min prior to treatment. Four different treatments (all at room temperature) were applied prior to 10-min HS at 55°C: (i) 10 mM EGTA, pH 8.0 for 10 min, (ii) 0.1% DMSO for 2 h, (iii) 15 μM CsA for 2 h, and (iv) 10 mM EGTA (applied 10 min prior to HS) and 15 μM CsA (applied 2 h prior to HS). Treatment times were staggered and samples were scanned within 10 min following HS. Cells were mounted on a microscope slide and imaged using Zeiss LSM 800.

In order to assess the cell death rate following EGTA treatment, an additional experiment was carried out using BY-2 cells that were treated and then stained with SO and FM4-64. There were three biological replicates for each treatment group: (i) No HS, control (water), (ii) No HS, with EGTA, (iii) 10-min 55°C HS, control, and (iv) 10-min 55°C HS, with EGTA. After HS, 300 μL of each sample was transferred to a 24-well plate designed for fluorescent microscopy (μ-Plate 24, Ibidi 82406) and imaged using Zeiss LSM 780 as described in the *Confocal microscopy* section.

### ATP content measurement

The assay was performed based on the previously published protocol (110). Half a mL aliquots of a 4-day-old BY-2 cell culture were transferred with a cut tip into 2 mL Eppendorf tubes. Six replicates were measured for each treatment in each experiment. The aliquoted cells were subjected to the treatments described in the *Fluorescein leakage* assay. Additionally, six aliquots were incubated with 48 μM CCCP on the bench for 1 h. After treatment, cells were pelleted on a nylon mesh with 50 μm pores, rinsed with 3 mL of medium and washed off into new Eppendorf tubes with 600 μL of boiling buffer (100 mM Tris-HCl, 4 mM EDTA, pH 7.5, autoclaved and supplemented with phosphatase inhibitor cocktail 2 (Sigma-Aldrich, P5726)). During sample collection, the tubes were kept on ice. Samples were then boiled at 100°C for 10 min, and debris was spun down at 4°C, 10,000*g* for 20 min. The supernatants were transferred into new tubes and stored on ice.

For each sample, 83.5 μL of supernatant were transferred into wells of white Nunc plates in three technical replicates. The background signal was detected using FLUOstar Omega Microplate Reader (BMG LABTECH), with a lens filter taking ten measurements per well. 41.5 μL of freshly prepared reaction buffer (1.5 mM DTT, 100 μM Luciferin (Sigma L 6152), 5 μg/mL Luciferase from *Phontius pyralis* (Sigma L9506)) were pipetted into each well by the automated pump in a microplate reader. Luminescence was detected for each well after adding the reaction buffer and shaking the plate to ensure good mixing in the reaction volume. A serial dilution of ATP ranging from 1.8 10^−2^ M to 10^−8^ M was used to build a standard curve and calculate the amount of ATP in each sample. A one-way ANOVA with Dunnet’s test was performed using JMP 10, and the graph was built in Origin 2019.

### CCCP treatment

Half a mL of 4-5-day old BY-2 cell culture were transferred to 1.5 mL Eppendorf tubes using a cut 1 mL tip. Staining with SO and FM4-64 was performed as described above (protocols i and ii for no HS and HS treatment groups, respectively). The four treatment groups included: (i) 10-min treatment with 48 μM CCCP at room temperature followed by 55°C HS for 10 min, (ii) 0.1% DMSO (vehicle control) at room temperature for 10 min followed by 55° HS for 10 min, (iii) 10-min treatment with 48 μM CCCP and no HS, and (iv) 0.1% DMSO with no HS. The cultures were transferred to a 24-well plate designed for fluorescent microscopy (μ-Plate 24, Ibidi 82406). Scanning of the wells was performed automatically using CLSM, 10 minutes after treatment and then every 2 h for 24 h. Three replicates were carried out per treatment group.

A HS gradient experiment was also carried out, comparing control (DMSO) and CCCP-treated BY-2 cells. The cultures were stained and treated as per the CCCP experiment described above and then subjected to one of the following conditions: (i) no HS, (ii) 40°C, (iii) 45°C, (iv) 50°C, or (v) 55°C HS for 10 min. The cultures were scanned after 6 h and 24 h using CLSM. Three technical replicates were carried out per group.

### Osmotic stress assay

Half a mL of 4-5-day old BY-2 cell culture were transferred to 1.5 mL Eppendorf tubes using a cut 1 mL tip. Cells were stained with FDA and exposed to HS as described in the *Fluorescein leakage assay*. D-Mannitol was pre-measured and added to Eppendorf tubes so that the addition of 200 μL of culture would give a final concentration of 500 mM. After HS, the cultures were allowed to cool for 10 min at 25°C and then 200 μL was transferred to the tubes containing D-Mannitol. The cultures were mixed gently by pipetting to dissolve the D-Mannitol. Within 30 min of D-Mannitol treatment, the samples were mounted on sample glass and imaged using Zeiss LSM 800.

### Ferroptosis assays

Three mL of 4-5-day old BY-2 cell culture were transferred to a 6-well plate using a cut 1 mL tip. Five different treatments were applied to the cell cultures for 16 h during which time they were returned to the incubator and grown as described above: (i) 0.1% DMSO (vehicle control), (ii) 1 μM Fer-1 (Sigma SML0583), (iii) 10 μM Fer-1, (iv) 50 μM Fer-1 and (v) 100 μM Fer-1. After the 16-h treatments, 0.5 mL of culture from each well were transferred to 1.5 mL Eppendorf tubes using a cut 1 mL tip and separated into three treatments: (i) no HS, (ii) 55° HS for 10 min, or (iii) 85° HS for 10 min as described above. Following treatment, 10 μL of culture were transferred to 490 μL of fresh BY-2 media in 24-well plates designed for fluorescent microscopy (μ-Plate 24, Ibidi 82406). Staining with SO and FM4-64 was performed as described above (protocols i and iii for no HS and HS treatment groups, respectively). Three technical replicates were carried out for each group and the wells were scanned automatically using CLSM at the following intervals: 10 min, 3, 6, 9, 12, 15, 18, 21 and 24 h. Three independent experiments were carried out.

The effect of Fer-1 was also tested on *Arabidopsis thaliana* seedlings which were grown as described above. Six-day old seedlings were gently transferred to 6-well plates containing 3 mL of liquid half-strength MS medium and given one of the following three treatments for 16 h: (i) 0.1% DMSO (vehicle control), (ii) 1 μM Fer-1, and (iii) 10 μM Fer-1. The medium was pipetted gently over the seedlings to ensure that roots were submerged in the liquid prior to being placed back into the growth cabinet overnight. The following morning, the seedlings were carefully transferred to 1.5 mL Eppendorf tubes containing the same treatment media from the 6-well plates before subjection to one of the following three treatments: (i) no HS, (ii) 55° HS for 10 min, or (iii) 85° HS for 10 min as described above. The seedlings were then stained with SO and FM4-64 as described above (protocols i and iii for no HS and HS treatment groups, respectively) and placed back into the growth cabinet until scanning with CLSM at 3 h and 6 h post-HS. The early differentiation zone of the root was scanned for three seedlings per time point, and three independent experiments were carried out.

### Confocal microscopy

Micrographs were acquired using either a LSM 800 or LSM 780 confocal laser scanning microscope (Carl Zeiss) with GaAsP detectors. Micrographs were taken with four different objectives: x10 (NA0.45), x20 (NA0.8), x40 (NA1.2, water immersion), and x63 (NA1.2, water immersion). For Fig. 1A, B and supplementary video S1, z-stack acquisition with sequential scanning was performed with a x63 objective. Nuclear phenotyping using the BY-2 transgenic line (Fig. 1C) was done with z-stack acquisitions using a x20 objective. Fluorescein was excited at 488 nm and emission was detected from 499 – 560 nm. MitoTracker red was excited at 561 nm and emission was detected from 582 – 754 nm. SO was excited at 561 nm and emission was detected from 410 – 605 nm. FM4-64 excitation was 506 nm, the emission was detected from 650 – 700 nm. Images were acquired using ZEN blue software (version 2.5, Carl Zeiss) or Zen black (version 2.3).

## List of abbreviations

AL-PCD: apoptosis-like programmed cell death
AVD: apoptotic volume decrease
Bcl-2: B-cell lymphoma 2
BY-2: Bright Yellow-2 cells
CCCP: carbonyl cyanide m-chlorophenylhydrazone
CLSM: confocal laser scanning microscopy
CsA: cyclosporin A
DIC: differential interference contrast
EGTA: ethylene glycol-bis(2-aminoethylether)-N,N,N’,N’-tetraacetic acid
FDA: fluorescein diacetate
Fer-1: Ferrostatin-1
GFP: green fluorescent protein
HS: heat shock
IQR: interquartile range
MMP: mitochondrial membrane potential
MPT: mitochondrial permeability transition
MPTP: mitochondrial permeability transition pore
MS: Murashige and Skoog medium
NLR: Nod-like receptor
NLS: nuclear localization signal
PCD: programmed cell death
PM: plasma membrane
RCD: regulated cell death
SO: Sytox Orange

## Acknowledgements

We would like to thank Prof. S. Kamoun for the insightful comments to the preprint version of this work published on BioRxiv.

## Authors’ contributions

EAM, APS and PVB developed the concept of this study and are main contributors to writing the manuscript. EAM, AND and FB performed all experiments, analysed data and prepared figures. VG contributed to the discussions of the results. All authors read and approved the final manuscript.

## Availability of data and materials

Data generated and analyzed during this study are included in the published article and its supplementary information files.

## Additional files

**Additional file 1:**

**Video S1.** Shrinkage of plant protoplast caused by 55°C HS is not followed by fragmentation into apoptotic bodies. 3D reconstructions (maximum intensity projections) of FM4-64-stained BY-2 cells to visualize the PM. Staining was performed according to protocols i and ii for the control conditions (no HS) and HS treatments, respectively (see Methods, *Sytox Orange and FM4-64 staining*). The arrows indicate membrane structure vaguely resembling PM blebbing or apoptotic bodies, which were not observed in the control (no HS) sample. These structures are localized on the inner side of the PM and do not separate from the cell even at late stages of cell death. Scale bars, 20 μm.

**Additional file 2**

**Figure S1.** Time course analysis of gross morphological changes following 55°C HS. FM4-64 and SO dyes were used to visualize collapsed cells with permeabilized PM. After a 10-min HS at 55°C, BY-2 cells were stained and imaged at several time points ranging from 15 min to 72 h. Staining was performed as described in protocol iv (see Methods, *Sytox Orange and FM4-64 staining*). FM4-64 panels represent the corresponding boxed areas. Arrows indicate FM4-64 positive vesicles that were found within the boundaries of cell corpses. DIC, differential interference contrast. Scale bars, 20 μm.

**Figure S2.** HS-induced cell death is ATP- and Ca^2+^-independent process. Morphology of FDA-stained cells under normal conditions (No HS) and after a 55°C HS. Protoplast shrinkage is denoted by arrows. Pre-treatment with 48 μM CCCP for 10 min, 15 μM CsA for 2 h, or 10 mM EGTA for 10 min did not alleviate protoplast shrinkage upon HS. Each treatment was repeated at least twice. Scale bars, 20 μm.

