## Supplementary figures and images for "Apoptosis is not conserved in plants as revealed by critical examination of a model for plant apoptosis-like cell death"

### Additional File Figure S1

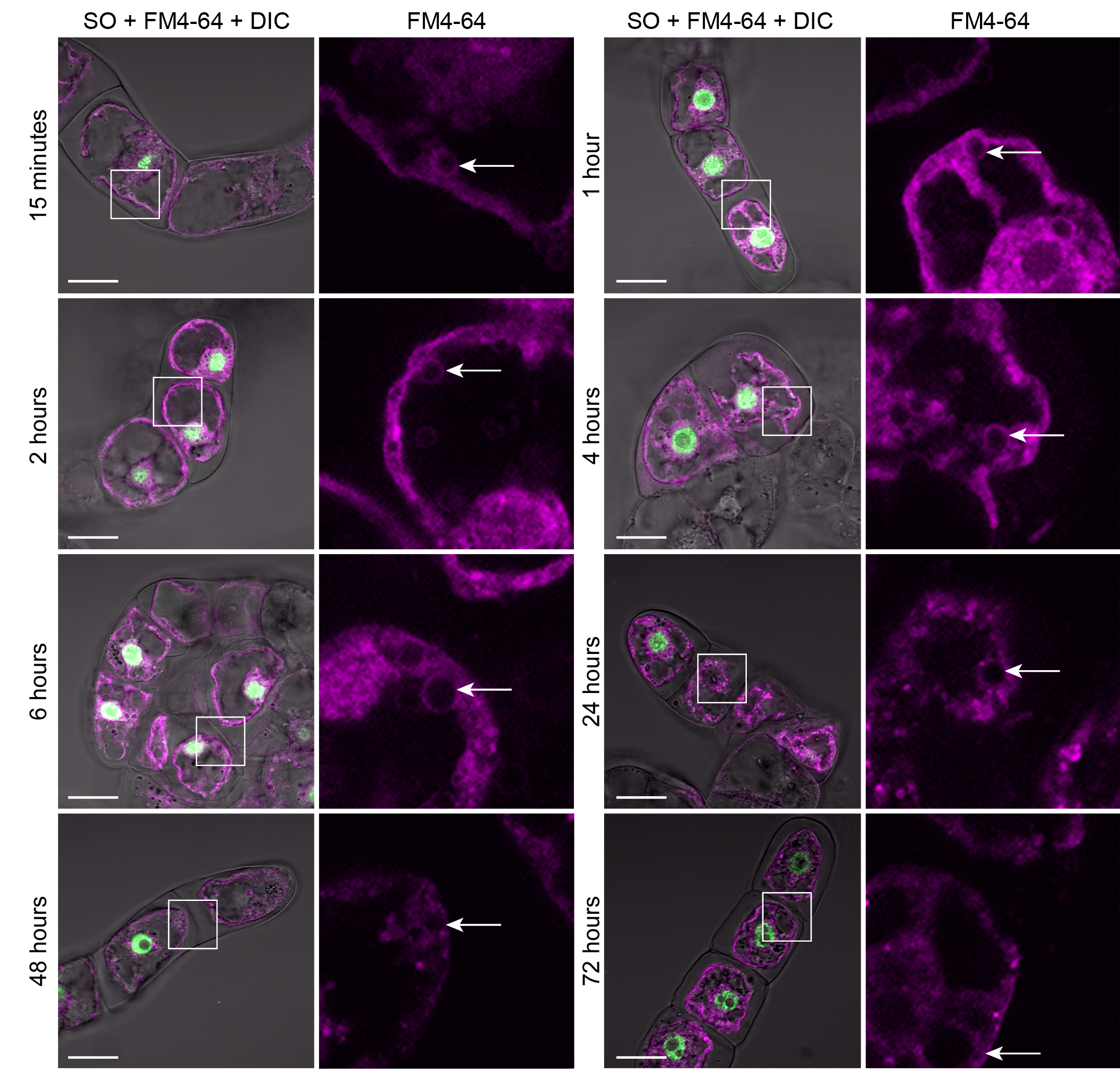

### Additional File Figure S2

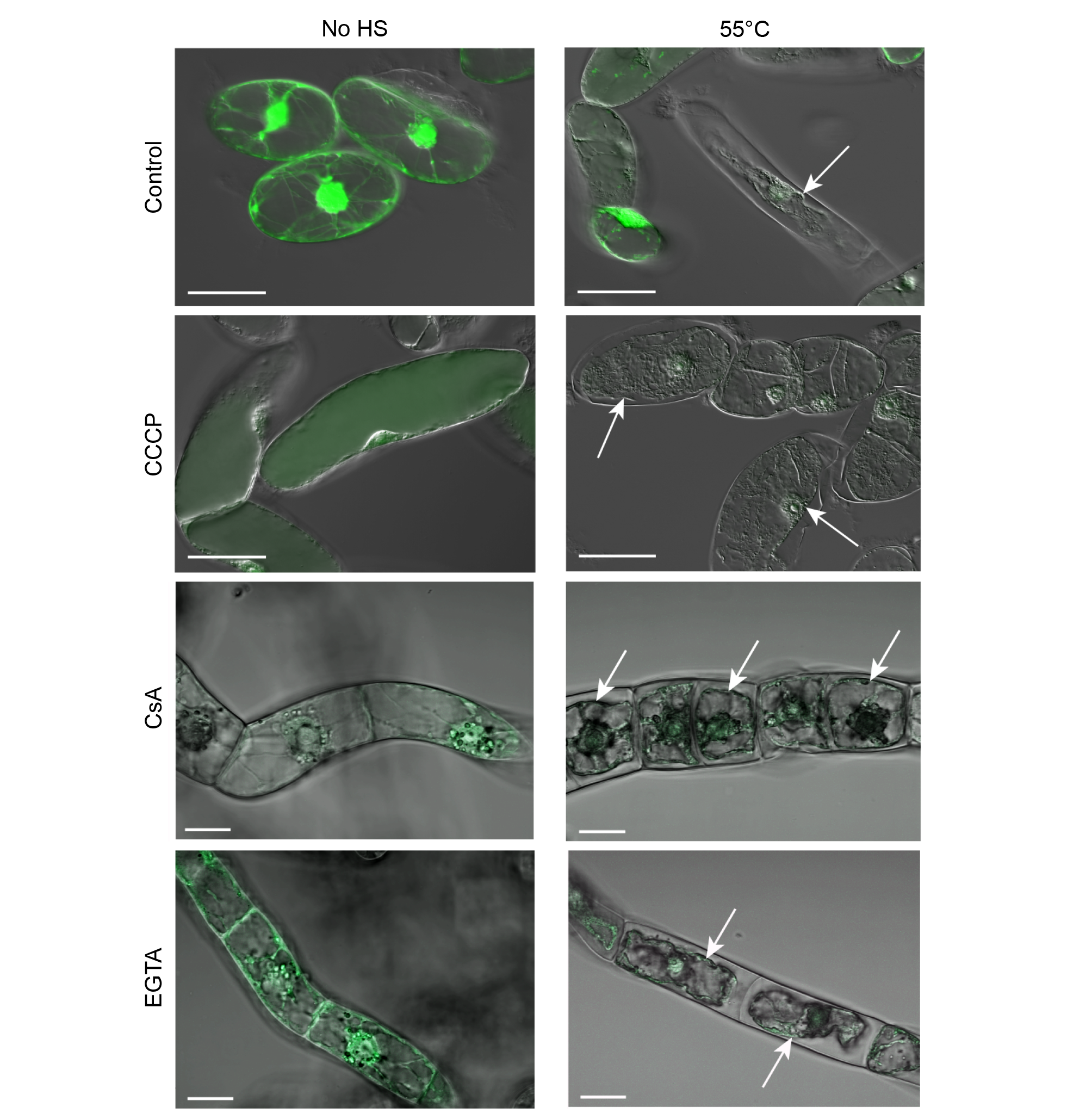
